# pysradb: A Python package to query next-generation sequencing metadata and data from NCBI Sequence Read Archive

**DOI:** 10.1101/578500

**Authors:** Saket Choudhary

**Affiliations:** Computational Biology and Bioinformatics, University Of Southern California, Los Angeles, CA 90089, USA

**Keywords:** bioinformatics, metadata, SRA, NGS, NCBI, GEO

## Abstract

NCBIs Sequence Read Archive (SRA) is the primary archive of next-generation sequencing datasets. SRA makes metadata and raw sequencing data available to the research community to encourage reproducibility, and to provide avenues for testing novel hypotheses on publicly available data. However, existing methods to programmatically access these data are limited. We introduce a Python package `pysradb` that provides a collection of command line methods to query and download metadata and data from SRA utilizing the curated metadata database available through the SRAdb project. We demonstrate the utility of `pysradb` on multiple use cases for searching and downloading SRA datasets. It is available freely at https://github.com/saketkc/pysradb.

## Introduction

Several projects have made efforts to analyze and publish summaries of DNA-seq [1] and RNA-seq [2, 3] datasets. Obtaining metadata and raw data from NCBI‘s Sequencing Read Archive (SRA) [4] is often the first step towards re-analyzing public next-generation sequencing datasets to compare them to private data or test a novel hypothesis. NCBI‘s SRA toolkit [5] provides utility methods to download raw sequencing data, while the metadata can be obtained by querying the website or through the Entrez `efetch` command line utility [6]. Most workflows analyzing public data rely on first searching for relevant keywords in the metadata either through the command line utility or the website, gathering relevant sample (s) of interest and then downloading these. A more streamlined workflow can enable performing all these steps at once.

In order to make querying both metadata and data more precise and robust, SRAdb [7] project provides a frequently updated SQLite database containing all the metadata parsed from SRA. `SRAdb` tracks the five main data objects in SRA‘s metadata: submission, study, sample, experiment and run. These are mapped to five different relational database tables that are made available in the SQLite file. The metadata semantics in the file remain as they are on SRA. The accompanying package `SRAdb` [8] made available in the R programming language [9] provides a convenient framework to handle metadata query and raw data downloads by utilizing the SQLite database. Though powerful, SRAdb requires the end user to be familiar with the R programming language and does not provide a command-line interface for querying or downloading operations.

`pysradb` package builds upon the principles of `SRAdb` providing a simple and user-friendly command-line interface for querying metadata and downloading datasets from SRA. It obviates the need for the user to be familiar with any programming language for querying and downloading datasets from SRA. Additionally, it provides utility functions that will further help a user perform more granular queries, that are often required when dealing with multiple datasets at large scale. By enabling both metadata search and download operations at the command-line, `pysradb` aims to bridge the gap in seamlessly retrieving public sequencing datasets and the associated metadata.

`pysradb` is written in Python (Python Software Foundation, https://www.python.org/) [10] and is currently developed on Github under the open-source BSD 3-Clause License. In order to simplify the installation procedure for the end-user, it is also available for download through PyPI (https://pypi.org/project/pysradb) and bioconda [11] (https://bioconda.github.io/recipes/pysradb/README.html)

## Methods

### Implementation

`pysradb` is implemented in Python and uses `pandas` [12] for data frame based operations. Since, downloading datasets can often take long time, `pysradb` displays progress for long haul tasks using `tqdm` [13]. The metadata information is read in the form of a SQLite [14] database made available by SRAdb [7].

`pysradb` can be run on either Linux or Mac based operating systems. It has minimal dependencies and can be easily installed using either `pip` or `conda` based package manager via the `bioconda` [11] channel.

Each sub-command of `pysradb` contains a self-contained help string, that describes its purpose and usage example. The help text can be accessed by passing the ‘–help’ flag. There is also additional documentation available for the sub-commands on the project‘s website (https://saketkc.github.io/pysradb). We also provide example Jupyter [15] notebooks (https://github.com/saketkc/pysradb/tree/master/notebooks) that demonstrate the functionality of the Python API.

`pysradb`’s development primarily happen on Github and the code is tested continuously using Travis CI webhook. This monitors all incoming pull requests and commits to the master branch. The testing happens on Python version 3.5, 3.6, and 3.7 on an Ubuntu 16.04 LTS virtual machine while the testing webhooks on the bioconda channel provides additional testing on Mac based systems.

## Use Cases

`pysradb` provides a chain of sub-commands for retrieving metadata, converting one accession to another and downloading. Each sub-command is designed to perform a single operation by default while additional operations can be performed by passing additional flags. In the following section we demonstrate some of the use cases of these sub-commands.

`pysradb` uses `SRAmetadb.sqlite`, a SQLite file produced and made available by SRAdb [7] project. The file itself can be downloaded using `pysradb` as:

`$ pysradb srametadb`

`SRAmetadb.sqlite` file is required for all other operations supported by `pysradb`. This file is required for all the sub-commands to function By default, `pysradb` assumes that by default the file is located in the current working directory. Alternatively, it can supplied as ‘–db path/to/SRAmetadb.sqlite’ argument. The examples here were run using `SRAmetadb.sqlite` with schema version 1.0 and creation timestamp 2019-01-25 00:38:19.

### Search

Consider a case where a user is looking for Ribo-seq [16] public datasets on SRA. These datasets will often have ‘ribosome profiling’ appearing in the abstract or sample description. We can search for such projects using the ‘search’ sub-command:

`$ pysradb search ‘"ribosome profiling"’ ∣ head`

| study_accession | experiment_accession | sample_accession | run_accession |
| --- | --- | --- | --- |
| DRP003075 | DRX019536 | DRS026974 | DRR021383 |
| DRP003075 | DRX019537 | DRS026982 | DRR021384 |
| DRP003075 | DRX019538 | DRS026979 | DRR021385 |
| DRP003075 | DRX019540 | DRS026984 | DRR021387 |
| DRP003075 | DRX019541 | DRS026978 | DRR021388 |
| DRP003075 | DRX019543 | DRS026980 | DRR021390 |
| DRP003075 | DRX019544 | DRS026981 | DRR021391 |
| ERP013565 | ERX1264364 | ERS1016056 | ERR1190989 |

The result here lists down all relevant ‘ribosome profiling’ projects.

### Getting metadata for a project (SRP)

Each SRA project (SRP) on NCBI’s SRA consists of one or multiple experiments (SRX) which are sequenced as one or multiple runs (SRR). Each experiment in turn is carried on a biological sample (SRS).

`pysradb metadata` can obtain all the experiment, sample, and run accessions associated with a project (SRP) as:

`$ pysradb metadata SRP098789 ∣ head`

| study_accession | experiment_accession | sample_accession | run_accession |
| --- | --- | --- | --- |
| SRP010679 | SRX118285 | SRS290854 | SRR403882 |
| SRP010679 | SRX118286 | SRS290855 | SRR403883 |
| SRP010679 | SRX118287 | SRS290856 | SRR403884 |
| SRP010679 | SRX118288 | SRS290857 | SRR403885 |
| SRP010679 | SRX118289 | SRS290858 | SRR403886 |
| SRP010679 | SRX118290 | SRS290859 | SRR403887 |
| SRP010679 | SRX118291 | SRS290860 | SRR403888 |
| SRP010679 | SRX118292 | SRS290861 | SRR403889 |
| SRP010679 | SRX118293 | SRS290862 | SRR403890 |
| SRP010679 | SRX118294 | SRS290863 | SRR403891 |
| SRP010679 | SRX118295 | SRS290864 | SRR403892 |
| SRP010679 | SRX118296 | SRS290865 | SRR403893 |

However, this information by itself is often not enough. We require detailed metadata associated with each sample to perform any downstream analysis. For example, the assays used for different samples and the corresponding treatment conditions. This can be done by supplying the ‘–desc’ flag:

`$ pysradb metadata SRP010679 –desc ∣ head −5`

|  |  |  |  |  |
| --- | --- | --- | --- | --- |
| study_accession | experiment_accession | sample_accession | run_accession | sample_attribute |
| SRP010679 | SRX118285 | SRS290854 | SRR403882 | source_name: PC3<br>human prostate<br>cancer cells \|\| cell<br>line: PC3 \|\| sample<br>type: polyA RNA \|\|<br>treatment: vehicle |
| SRP010679 | SRX118286 | SRS290855 | SRR403883 | source_name: PC3<br>human prostate cancer<br>cells \|\| cell line:<br>PC3 \|\| sample type:<br>ribosome protected<br>RNA \|\| treatment:<br>vehicle |
| SRP010679 | SRX118287 | SRS290856 | SRR403884 | source_name: PC3<br>human prostate<br>cancer cells \|\| cell<br>line: PC3 \|\| sample<br>type: polyA RNA \|\|<br>treatment: rapamycin |
| SRP010679 | SRX118288 | SRS290857 | SRR403885 | source_name: PC3<br>human prostate cancer<br>cells \|\| cell line:<br>PC3 \|\| sample type:<br>ribosome protected<br>RNA \|\| treatment:<br>rapamycin |

This can be further expanded to reveal the data in ‘sample_attribute’ column into separate columns via ‘–expand’ flag. This is most useful for samples that have treatment or cell type associated metadata available.

`$ pysradb metadata SRP010679 –desc –expand`

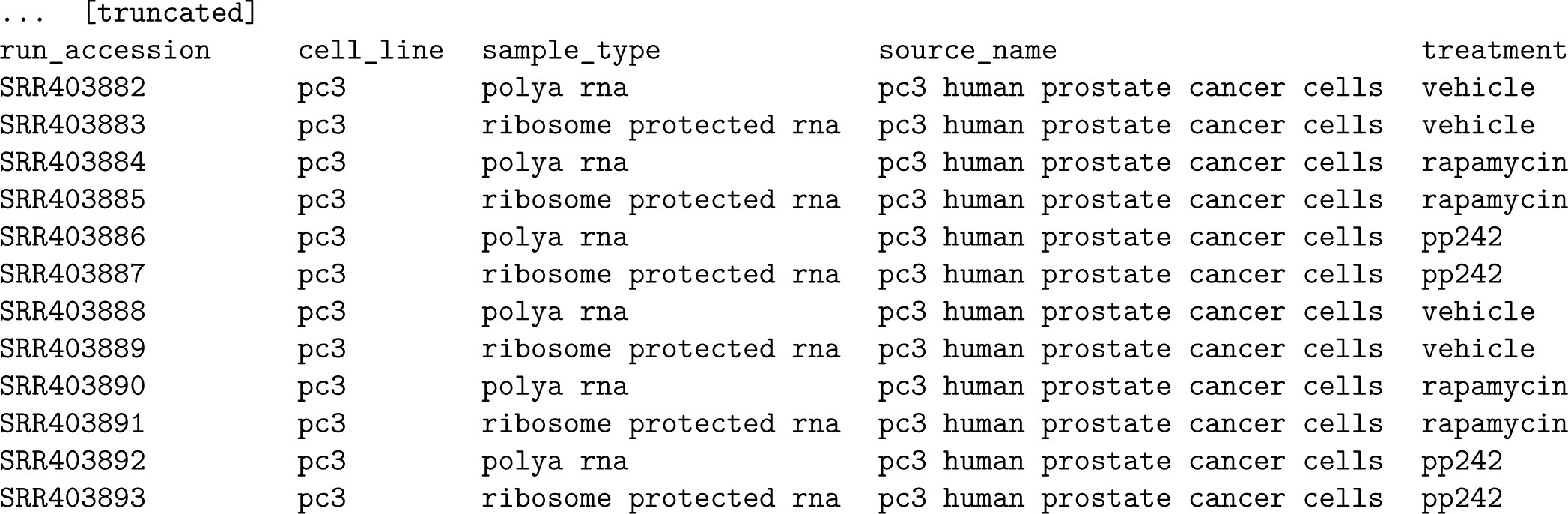

Any SRA project might consist of experiments involving multiple assay types. The assay associated with any project can be obtained by providing `–assay` flag:

`$ pysradb metadata SRP000941 –assay ∣ tr -s ‘ ’ ∣ cut -f5 -d ‘ ’ ∣ tail -n +2 ∣ sort ∣ uniq -c`

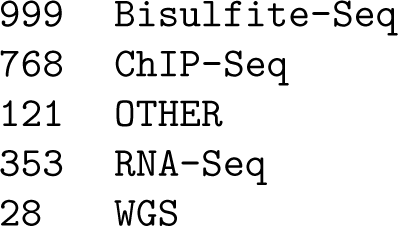

### Getting SRPs from GSE

The Gene Expression Omnibus database (GEO) [17] is NCBI’s data repository for functional genomics data that accepts array and sequence-based data from gene profiling experiments. For sequence-data, the corresponding raw files are deposited to the SRA. GEO assigns its datasets accession (GSE) that are linked to their corresponding accession on the SRA (SRP). It is often required to interpolate between the two accessions. The `gse-to-srp` sub-command allows converting GSE to SRP:

`$ pysradb gse-to-srp GSE24355 GSE25842`

| study_alias | study_accession |
| --- | --- |
| GSE24355 | SRP003870 |
| GSE25842 | SRP005378 |

It can further be expanded to obtain the corresponding experiment and run accessions:

`$ pysradb gse-to-srp –detailed –expand GSE100007 ∣ head`

| study_alias | study_accession | experiment_accession | sample_accession | experiment_alias | sample_alias |
| --- | --- | --- | --- | --- | --- |
| GSE100007 | SRP109126 | SRX2916198 | SRS2282390 | GSM2667747 | GSM2667747 |
| GSE100007 | SRP109126 | SRX2916199 | SRS2282391 | GSM2667748 | GSM2667748 |
| GSE100007 | SRP109126 | SRX2916200 | SRS2282392 | GSM2667749 | GSM2667749 |
| GSE100007 | SRP109126 | SRX2916201 | SRS2282393 | GSM2667750 | GSM2667750 |
| GSE100007 | SRP109126 | SRX2916202 | SRS2282394 | GSM2667751 | GSM2667751 |
| GSE100007 | SRP109126 | SRX2916203 | SRS2282395 | GSM2667752 | GSM2667752 |
| GSE100007 | SRP109126 | SRX2916204 | SRS2282396 | GSM2667753 | GSM2667753 |
| GSE100007 | SRP109126 | SRX2916205 | SRS2282397 | GSM2667754 | GSM2667754 |
| GSE100007 | SRP109126 | SRX2916206 | SRS2282400 | GSM2667755 | GSM2667755 |

### Getting a list of GEO experiments for a GEO study

For any GEO study (GSE) that involves a collection of experiments (GSM), we can obtain an entire list of experiments using the `gse-to-gsm` sub-command from `pysradb`:

`$ pysradb gse-to-gsm GSE41637 ∣ head`

| study_alias | experiment_alias |
| --- | --- |
| GSE41637 | GSM1020640_1 |
| GSE41637 | GSM1020641_1 |
| GSE41637 | GSM1020642_1 |
| GSE41637 | GSM1020643_1 |
| GSE41637 | GSM1020644_1 |
| GSE41637 | GSM1020645_1 |
| GSE41637 | GSM1020646_1 |
| GSE41637 | GSM1020647_1 |
| GSE41637 | GSM1020648_1 |

However, just a list of GSM accessions is not useful if one is performing any downstream analysis which essentially requires more detailed information about the metadata associated with each experiment. This relevant metadata associated with each sample can be obtained by providing `gse-to-gsm` additional flags:

`$ pysradb gse-to-gsm –desc GSE41637 ∣ head`

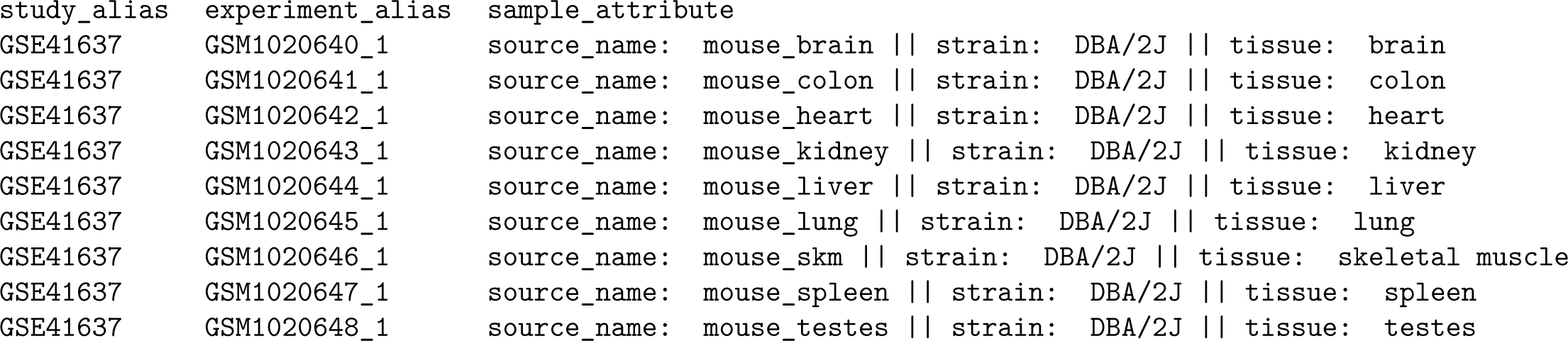

| study_alias | experiment_alias | sample_attribute |
| --- | --- | --- |
| GSE41637 | GSM1020640_1 | source_name: mouse_brain \|\| strain: DBA/2J \|\| tissue: brain |
| GSE41637 | GSM1020641_1 | source_name: mouse_colon \|\| strain: DBA/2J \|\| tissue: colon |
| GSE41637 | GSM1020642_1 | source_name: mouse_heart \|\| strain: DBA/2J \|\| tissue: heart |
| GSE41637 | GSM1020643_1 | source_name: mouse_kidney \|\| strain: DBA/2J \|\| tissue: kidney |
| GSE41637 | GSM1020644_1 | source_name: mouse_liver \|\| strain: DBA/2J \|\| tissue: liver |
| GSE41637 | GSM1020645_1 | source_name: mouse_lung \|\| strain: DBA/2J \|\| tissue: lung |
| GSE41637 | GSM1020646_1 | source_name: mouse_skm \|\| strain: DBA/2J \|\| tissue: skeletal muscle |
| GSE41637 | GSM1020647_1 | source_name: mouse_spleen \|\| strain: DBA/2J \|\| tissue: spleen |
| GSE41637 | GSM1020648_1 | source_name: mouse_testes \|\| strain: DBA/2J \|\| tissue: testes |

The metadata information can then be parsed from the `sample_attribute` column. To obtain more structured metadata, we can use an additional flag ‘`–expand`’:

`$ pysradb gse-to-gsm –desc –expand GSE41637 ∣ head`

| study_alias | experiment_alias | source_name | strain | tissue |
| --- | --- | --- | --- | --- |
| GSE41637 | GSM1020640_1 | mouse_brain | dba/2j | brain |
| GSE41637 | GSM1020641_1 | mouse_colon | dba/2j | colon |
| GSE41637 | GSM1020642_1 | mouse_heart | dba/2j | heart |
| GSE41637 | GSM1020643_1 | mouse_kidney | dba/2j | kidney |
| GSE41637 | GSM1020644_1 | mouse_liver | dba/2j | liver |
| GSE41637 | GSM1020645_1 | mouse_lung | dba/2j | lung |
| GSE41637 | GSM1020646_1 | mouse_skm | dba/2j | skeletal muscle |

### Getting SRR from GSM

`gsm-to-srr` allows conversion from GEO experiments (GSM) to SRA runs (SRR):

`$ pysradb gsm-to-srr GSM1020640 GSM1020646`

| experiment_alias | run_accession |
| --- | --- |
| GSM1020640_1 | SRR594393 |
| GSM1020646_1 | SRR594399 |

### Downloading SRA datasets

`pysradb` enables seemless downloads from NCBI’s SRA. It organizes the downloaded data following NCBI‘s hiererachy: ‘SRP => SRX => SRR’ of storing data. Each ‘SRP’ (project) has multiple ‘SRX’ (experiments) and each ‘SRX’ in turn has multiple ‘SRR’ (runs). Multiple projects can be downloaded at once using the `download` sub-command:

`$ pysradb download -p SRP003870 -p SRP005378`

`download` also allows Unix pipes based inputs. Consider our previous example of the project SRP000941 with different assays. However, we want to be able to download only ‘RNA-seq’ samples. We can do this by subsetting the metadata output for only ‘RNA-seq’ samples:

`$ pysradb metadata SRP000941 –assay ∣ grep ‘study∣RNA-Seq’ ∣ pysradb download`

This will only download the ‘RNA-seq’ samples from the project.

## Summary

`pysradb` provides a command-line interface to query metadata and download sequencing datasets from NCBI‘s SRA. It enables seamless retrieval of metadata and conversion between different accessions. `pysradb` is written in Python 3 and is available on Linux and Mac OS. The source code is hosted on Github and licensed under BSD 3-clause license. It is available for installation through PyPI and bioconda.

## Software availability

Software and source code available from: https://github.com/saketkc/pysradb

Documentation available at: https://saketkc.github.io/pysradb

Archived source code at time of publication: https://doi.org/10.5281/zenodo.2579446

Software license: BSD 3-Clause

## Author Contributions

S.C. designed the project, implemented the package, and wrote the manuscript.

## Acknowledgments

The author thanks Amal Thomas, Meng Zhou, Rishvanth Prabakar, Wenzheng Li, and Xiaojing Ji at the University of Southern California (USC) and Shweta Ramdas at the University of Pennsylvania for helpful discussions and comments on the software and manuscript. The author acknowledges support from the USC Provost Graduate Research Fellowship.

